# Bayesian inference of ancestral recombination graphs for bacterial populations

**DOI:** 10.1101/059105

**Authors:** Timothy G. Vaughan, David Welch, Alexei J. Drummond, Patrick J. Biggs, Tessy George, Nigel P. French

**Affiliations:** Centre for Computational Evolution and Department of Computer Science, University of Auckland, Auckland, New Zealand; mEpiLab, Infectious Disease Research Centre, Hopkirk Research Institute, Massey University, Palmerston North, New Zealand

**Keywords:** bacterial evolution, recombination, phylogenetic inference

## Abstract

Homologous recombination is a central feature of bacterial evolution, yet confounds traditional phylogenetic methods. While a number of methods specific to bacterial evolution have been developed, none of these permit joint inference of a bacterial recombination graph and associated parameters. In this paper, we present a new method which addresses this shortcoming. Our method uses a novel Markov chain Monte Carlo algorithm to perform phylogenetic inference under the ClonalOrigin model of Didelot et al. (Genetics, 2010). We demonstrate the utility of our method by applying it to rMLST data sequenced from pathogenic and non-pathogenic *Escherichia coli* serotype O157 and O26 isolates collected in rural New Zealand. The method is implemented as an open source BEAST 2 package, Bacter, which is available via the project web page at tgvaughan.github.io/bacter

---

RECOMBINATION plays a crucial role in the molecular evolution of many bacteria, in spite of the clonal nature of bacterial reproduction. Indeed, for a large number of species surveyed in recent studies (Vos and Didelot 2009; Fearnhead *et al.* 2015), homologous recombination was found to account for a similar or greater number of nucleotide changes than point mutation.

Yet many traditional phylogenetic methods (Huelsenbeck and Ronquist 2001; Drummond et *al.* 2002; Guindon and Gascuel 2003) do not account for recombination. This is regrettable for several reasons. First, ignoring recombination is known to bias phylogenetic analyses in various ways such as by overestimating the number of mutations along branches, artificially degrading the molecular clock hypothesis, and introducing apparent exponential population growth (Schierup and Hein 2000). Second, much of modern computational phylogenetics extends beyond the inference of phylogenetic relationships and instead focuses on the parametric and non-parametric inference of the dynamics governing the population from which the genetic data is sampled. In this context, the phylogeny is merely the glue that ties the data to the underlying population dynamics. Recombination events contain a strong phylogenetic signal so incorporating recombination into the phylogenetic model can significantly improve analyses. For instance, Li and Durbin (2011) used a recombination-aware model to recover detailed ancestral population dynamics from pairs of human autosomes, a feat which would have been impossible without the additional signal provided by the recombination process.

The standard representation of the phylogenetic relationship between ancestral lineages when recombination is present is the Ancestral Recombination Graph (ARG Griffiths 1981; Hudson 1983), a timed phylogenetic network describing the reticulated ancestry of a set of sampled taxa. Several inference methods based on the ARG concept have been developed, many of which (Wang and Rannala 2008; Bloomquist and Suchard 2010; Li and Durbin 2011) assume a symmetry between the contributions of genetic material from the parent individuals contributing to each recombination event, as is the expected result of the crossover resolution of the Holliday junction in eukaryotic recombination. This assumption, which is often anchored in the choice to base the inference on the coalescent with recombination (Wiuf and Hein 1999), is not generally appropriate for bacterial recombination, where there is usually a strong asymmetry between the quantity of genetic material contributed from each ‘parent’.

Alternatively, a series of methods introduced by Didelot and coauthors (Didelot and Falush 2007; Didelot et *al.* 2010; Didelot and Wilson 2015) directly target bacterial recombination by employing models based on the coalescent with gene conversion (Hudson 1983; Wiuf 2000; Wiuf and Hein 2000). These models acknowledge that the asymmetry present in the bacterial context allows for the definition of a precisely defined clonal genealogy-the *clonal frame*-which represents not only the true reproductive genealogy of a given set of bacterial samples, but also the ancestry of the majority of their genetic material.

In the first paper, Didelot and Falush (2007) presented a method for performing inference under a model of molecular evolution which, in combination with a standard substitution model, includes effects similar to those resulting from gene conversion; instantaneous events which simultaneously produce character state changes at multiple sites within a randomly positioned conversion tract. This model does not consider the origin of these changes: it dispenses entirely with the ARG and can be considered a rather peculiar substitution model applied to evolution of sequences down the clonal frame. Despite this, it does allow the MCMC algorithm implemented in the associated ClonalFrame software package to jointly infer the bacterial clonal frame, conversion rate and tract length parameters, neatly avoiding the branch length bias described by Schierup and Hein (2000). Didelot and Wilson (2015) introduced a maximum likelihood method for performing inference under the same model, making it possible to infer clonal frames from whole bacterial genomes as opposed to the short seqeunces that the earlier Bayesian method could handle.

In a second paper, Didelot *et al.* (2010) present a different approximation to the coalescent with gene conversion which retains the ARG but assumes that the ARG has the form of a tree-based network (Zhang 2015) with the clonal frame taking on the role of the base tree. While acknowledging that their model could be applied to jointly infer the clonal frame and the conversions, the algorithm they present is limited to performing inference of the gene conversion ARG given a separately-inferred clonal frame. This choice permitted the application of their model to relatively large genomic data sets.

This model was also used recently by Ansari and Didelot (2014), who exploit the Markov property of the model with regard to the active conversions at each site along an aligned set of sequences to enable rapid simulation under the model. These simulations were used in an approximate Bayesian computation scheme (Beaumont *et al.* 2002) to infer the homologous recombination rate, tract lengths and scaled mutation rate from full genome data, as well as to assess the degree to which the recombination process favours DNA from donors closely related to the recipient. As with the earlier study, this method requires that the clonal frame be separately inferred.

In this paper we present a Bayesian method for jointly reconstructing the ARG, the homologous conversion events, the expected conversion rate and tract lengths and the population history from genetic sequence data. Our approach assumes the ClonalOrigin model of Didelot *et al.* (2010), extended to allow for the piecewise constant or piecewise linear variations in population size. It relies upon a novel Markov Chain Monte Carlo (MCMC) algorithm which uses a carefully designed set of proposal distributions in order to make traversing the vast state space of the model tractable for practical applications. Unlike earlier methods, our algorithm jointly infers the clonal frame, meaning that the inference is a single-step process.

In addition to the inference method itself, we present a basic technique for summarising the sampled ARG posterior. Our approach is an extension of the Maximum Clade Credibility tree approach (as described by Heled and Bouckaert 2013) to summarising phylogenetic tree posteriors in which a summary of the clonal frame is annotated with well-supported conversion events.

We demonstrate that our method can accurately infer known parameters from simulated data and apply it to a set of *Escherichia coli* rMLST (ribosomal multi-locus sequence typing, Jolley *et al.* 2012) sequences derived from isolates collected from in and around the Manawatu region in New Zealand. The method reveals details of previously unobserved gene flow between pathogenic and non-pathogenic populations belonging to the serotype O157.

A software implementation of our method is distributed as a publicly-available BEAST 2 (Bouckaert *et al.* 2014) package. This gives the sampler a substantial amount of flexibility, allowing it to be used in combination with complex substitution models and a wide variety of prior distributions. Details on how to obtain and use this package are given on the project website at tgvaughan.github.io/bacter

## The ClonalOrigin genealogical model

In contrast to eukaryotes where recombination primarily occurs during meiosis, bacteria generally undergo recombination due to mechanisms that are not directly related to the process of genome replication. These mechanisms generally only result in the transfer of small fragments of genetic material. A result of this is that every homologous recombination event in bacteria is comparable to a gene conversion event, regardless of the underlying molecular biology. A good model for the genealogy of bacterial genomes is therefore the coalescent with gene conversion: a straight-forward extension to the Kingman *n*-coalescent (Kingman 1982a,b) in which (a) lineages may bifurcate as well as coalesce and (b) lineages are associated with a subset of sites on each of the sampled genetic sequences to which they are ancestral. At each bifurcation event, a contiguous range of sites is chosen for “conversion” by selecting a starting site uniformly at random and a tract length from a geometric distribution. The ancestry of the converted sites follows one parental lineage, while that of the unconverted sites follows the other.

The ClonalOrigin model is a simplification of the coalescent with gene conversion in which lineages are labeled as either clonal or non-clonal, with non-clonal lineages assumed to be free from conversion events (i.e. they may not bifurcate) and pairs of these lineages forbidden from coalescing. As Didelot *et al.* (2010) argue, this simplified process is a good approximation to the full model in the limit of small expected tract length (relative to genome length) and low recombination rate. It also possesses features that make it an attractive basis for practical inference methods. First among these is that, conditional on the clonal frame (CF), the conversion events are completely independent. In our context, this simplifies the process of computing the probability of a given ARG and proposing the modifications necessary when exploring ARG-space using MCMC.

We briefly reiterate the mathematical details of the model described in Didelot *et al.* (2010) using terminology more appropriate for our purposes. We define the ClonalOrigin recombination graph *G* = (*C*, *R*) where *C* represents the clonal frame and *R* is a set of recombinant edges connecting pairs of points on C. The CF is assumed to be generated by an unstructured coalescent process governed by a time-dependent effective population size *N*(*t*), where *t* measures time before the present. That is, the probability density of *C* can be written

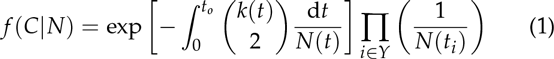

Here *Y* is the set of internal (coalescent) nodes between edges of *C*, including the root node o, and {*t*_*j*_|*i* ϵ *Y*} are the ages of these nodes. The term *k*(*t*) represents the number of CF lineages extant at time *t*.

**Figure 1.**
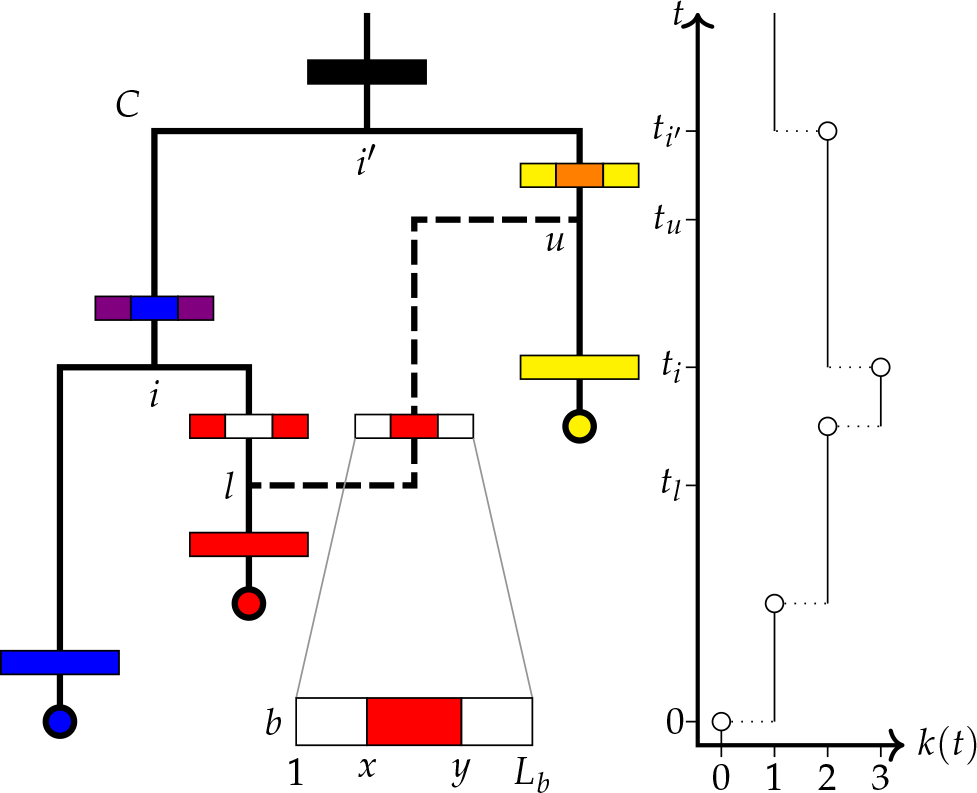
Schematic representation of a recombination graph *G* for a single locus *b*, with clonal frame *C* and |*R*| = 1 conversion *r*. The conversion attaches to *C* at points *l* and *u* and affects sites *x* through *y* of the *L_b_* sites belonging to locus *b*. The horizontal bars represent ancestral sequences belonging to each lineage and colors are used to denote which samples each site is ancestral to, with white indicating sites ancestral to no samples. The graph on the right displays the associated clonal frame lineages-through-time function *k*(*t*), together with the times used in computing *f* (*G*|*N*, *δ*, *ρ*, *B*). These include the conversion attachment times *t_l_*; and *t_u_*, together with ages of coalescent nodes *i* and *i*′. (Here *i*′ = *o*.)

Conversion events *r* ϵ *R* appear at a constant rate on each lineage of *C* and thus their number |*R*| is Poisson-distributed with mean *T*Σ_*b*ϵ*B*_(*ρL*_*b*_+ *δ* − 1), with *T* being the sum of all branch lengths in *C*. Here *ρ* is the per site per unit time rate of homologous gene conversion, *δ* is the expected conversion tract length and *b* ϵ *B* are the loci for which length *L_b_* sequence alignments are available. Each conversion is defined by *r* = (*l, u, b, x, y*) where *l* and *u* identify points on *C* at which the recombinant lineage attaches, with the age of *l* less than that of *u*. The element *b* indicates the locus to which the conversion applies, and *x* and *y* identify the start and end respectively of the range of sites affected by the conversion. The point *l* ~ *f* (*l*|*C*) is chosen uniformly over *C*, while *u* is drawn from the conditional coalescent distribution
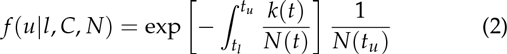
where *t*_*l*_ and *t*_*u*_ are the ages of points *l* and *u* respectively. The locus *b* is chosen with probability *P*(*b*|*B*,*δ*) = (*L_b_* + *δ* −1)/Σ_*b*′ ϵ *B*_(*L*_*b*′_ + *δ* − 1), the site *x* is drawn from the distribution *P*(*x*|*b*, *δ*) = (*I*(*x* = 1) / (*l* + *δ* −), and the site *y* is drawn from *P*(*y*|*x*, *b*, *δ*) = *δ*^−1^(1 − *δ*^−1^)^*y* − *x*^ + *I*(*y* = *L_b_*)(1 − *δ*^−1^)^*L_b_* − *x*^. (In these equations *I*(‧) is the indicator function.)

The full probability density for a ClonalOrigin ARG is then simply the product:
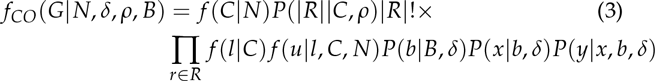
where the |*R*|! accounts for independence with respect to label permutations of the recombination set *R*. Figure 1 illustrates the various elements of the ClonalOrigin model and associated notation.

## Bayesian inference

Performing Bayesian inference under the ClonalOrigin model shares many similarities with the process of performing inference under the standard coalescent. The goal is to characterise the joint posterior density:
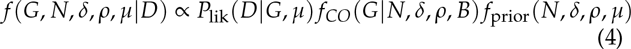
where *D* represents multiple sequence alignments for each locus in *B* and *μ* represents one or more parameters of the chosen substitution model. The distributions on the right-hand side include *P*_lik_, the likelihood of the recombination graph, *f_CO_*, the probability density of the graph under the ClonalOrigin model discussed above, and *f*_prior_, the joint prior density of the model parameters.

To define the ARG likelihood, first consider that every ARG may be mapped onto a set 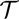 of “local” trees describing the ancestry of contiguous ranges of completely linked sites in the alignment. The likelihood of *G* is expressed in terms of local trees as the following product
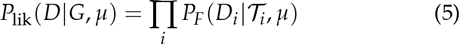
where *D_i_* is the portion of the alignment whose ancestry is described by local tree 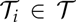 and 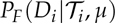 is the standard phylogenetic tree likelihood (Felsenstein 2003).

Since it is possible for conversions to have no effect on 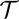, there is no one-to-one correspondence between *G* and 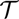. This suggests that certain features of *G* may be strictly non-identifiable in terms of the likelihood function. As Bayesian inference deals directly with the posterior distribution, this non-identifiability will not invalidate any analysis provided that f_prior_ is proper. However, the existence of non-identifiability has practical implications for the design of sampling algorithms, as we discuss in the following section.

### Markov chain Monte Carlo (MCMC)

We use MCMC to sample from the joint posterior given in eq. (4). This algorithm explores the state space of *x* = (*G*, *N*, *δ*, *ρ*, *μ*) (or some subspace thereof) using a random walk in which steps from *x* to *x*′ are drawn from some proposal distribution *q*(*x*′|*x*) and accepted with a probability that depends on the relative posterior densities at *x*′ and *x*.

In practice, *q*(*x*′|*x*) is often expressed as a weighted sum of proposal densities *q_i_* (*x*′|*x*) (also known as *proposals* or *moves*) which individually propose alterations to some small part of *x*. While there is considerable freedom in choosing a set of moves, their precise form can dramatically influence the convergence and efficiency of the sampling algorithm. Proposals should not generate new states that are too bold (accepted with very low frequency) nor too timid (accepted with very high frequency): both extremes tend to lead to chains with long autocorrelationperiods. In this section we present an informal outline of the moves used in our algorithm. (Refer to the Appendix for a detailed description.)

For the sub-space made up of the continuous model parameters *δ*, *ρ*, *μ*, and *N*, choosing appropriate proposals is relatively trivial as standard proposals for sampling from ℝ^*n*^ are sufficient. In our algorithm we use the univariate scaling operator described by Drummond *et al*. (2002), which can be made more or less bold simply by altering the size of the scaling operation.

For the ARG itself, assembling an appropriate set of moves is more difficult. Even determining exactly what constitutes a timid or bold move in G space is hard to determine without detailed knowledge of the target density. Our general approach here is to design proposals that (a) only minimally affect the likelihood of *G* where possible and (b) draw any significant changes from the prior that the ClonalOrigin model places on *G*. The design of these proposals is assisted by our knowledge of the identifiability issue considered in the previous section: there is a many-to-one mapping from *G* to the local tree set 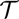, and the ARG likelihood depends only on 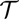. Thus, ARG proposals that minimally effect the likelihood are those that propose a *G′* that maps to the same or similar 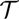.

The proposals for *G* fall into two groups, the first of which deals exclusively with the set of conversions *R*. These include all three moves described by Didelot *et al.* (2010) (we consider the conversion add/remove pair to be two halves of a single proposal), along with six additional simple moves aimed at quickly exploring the ARG state space conditional on *C*. Examples include a conversion merge/split proposal that merges pairs of conversions between the same pair of edges on the CF that affect nearby ranges of sites or splits single conversions into such pairs, a proposal which reversibly replaces a single conversion between two edges with a pair involving a third intermediate edge, and a proposal which adds or removes conversions that do not alter the topology of the CF.

Proposals in the second group propose joint updates to both the clonal frame *C* and the conversions *R*. Some of these moves are quite bold (and thus tend to be accepted rarely), but are very important for dealing with topological uncertainty in the clonal frame. The general strategy for each move is to apply one of the tree proposals from Drummond *et al.* (2002) to *C* and to simultaneously modify the conversions in *R* to ensure both compatibility with the *C*′ and to minimise the effect of the proposal on both the likelihood and the ARG prior. The changes to *C* can for the most part be decomposed into primitive operations that involve selecting a subtree, deleting the edge *e* attaching that subtree to the rest of the CF at time *t_i_*, then reconnecting the subtree via a new edge *e*′ to a new point on *C* at time *t*′_i_. Modification of *R* is done using an approach (depicted in figure 2) which consists of two distinct forms. The first form, the “collapse”, is applied whenever *t*′_*i*_ < *t_i_*; and involves finding conversions for which *u* or *v* are on the edge above the subtree and attach at times *t_l_* or *t_u_* greater than *t′_i_*. These attachment points are moved from their original position to contemporaneous points on the *C* lineage ancestral to *e*′. The second form, the “expansion”, is applied when *t′_i_* > *t_i_* and is the inverse of the first: conversion attachments *u* or *v* at times *t_i_* < *t_{l,u}_* < *t′_i_* are moved with some probability to contemporaneous positions on *e*′.

In concert, these proposals allow us to effectively explore the entire state space of *x*.

**Figure 2.**
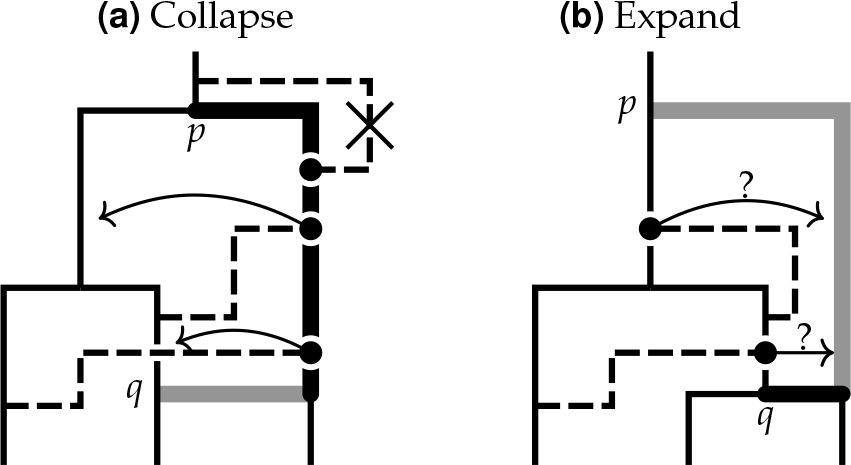
Schematic representation of the collapse/expand strategy used by the MCMC algorithm to update conversions following the movement of a clonal frame edge. Sub-figure (a) illustrates a proposal to replace the thick black edge portion of the clonal frame edge joined to *p* with the thick grey edge portion joint to *q*. Since *t_q_* < *t_p_* the “collapse” procedure is applied by moving affected conversion attachment points, highlighted with solid circles, to contemporaneous points on the lineage ancestral to *q*. Any conversion with a new arrival point above the root is deleted from the new ARG. Sub-figure
(b) illustrates the reverse situation, where a CF edge attached at q is reattached at *p*. Since *t_p_* > *t_q_* the “expand” procedure is applied by moving any attachment points contemporaneous with a point on the newly extended portion of the CF edge to that point with some probability. Since *p* becomes the new CF root, new conversions with arrival points on the new CF edge at times older than the previous CF root are drawn from the ClonalOrigin prior.

### A. Summarizing the ARG posterior

Bayesian MCMC algorithms produce samples from posterior distributions rather than point estimates of inferred quantities. These approaches are superior because they give us the means to directly quantify the uncertainty inherent in the inference. For the very high dimensional state space that ARGs (even the ClonalOrigin model′s tree-based networks) occupy, actually visualising this uncertainty and extracting an overall picture of the likely ancestral history of the sequence data is non-trivial.

A similar problem exists for Bayesian phylogenetic tree inference. Given the maturity of that field, it should not be surprising that a large number of solutions exist. The majority of these solutions involve the assembly of some kind of summary or consensus tree (see chapter 30 of Felsenstein (2003) for an overview or Heled and Bouckaert (2013) for a recent discussion). While conceptually appealing, the replacement of a posterior distribution with a single tree can very easily lead to the appearance of signal where there is none, so care must be taken. At least one method exists that avoids this problem: the DensiTree software (Bouckaert 2010) simply draws all of the trees in a given set with some degree of transparency, making it possible to actually visualize the distribution directly.

Unfortunately, the approach taken by DensiTree cannot be easily applied to ARGs, since the recombinant edges introduce significant visual noise making patterns difficult to discern. Nor can any of the standard summary methods be applied directly.

Instead, we use a summary of the CF posterior as a starting point to produce summary ARGs, as described in algorithm 1. In the algorithm, MCC refers to the Maximal Clade Credibility tree (see, for instance, Heled and Bouckaert 2013), and the value of *α* in step 3(c) imposes a threshold on the posterior support necessary for a conversion to appear in the summary. The relationship between the sampled conversions and the summary conversions is illustrated in figure 3.

**Algorithm 1.**
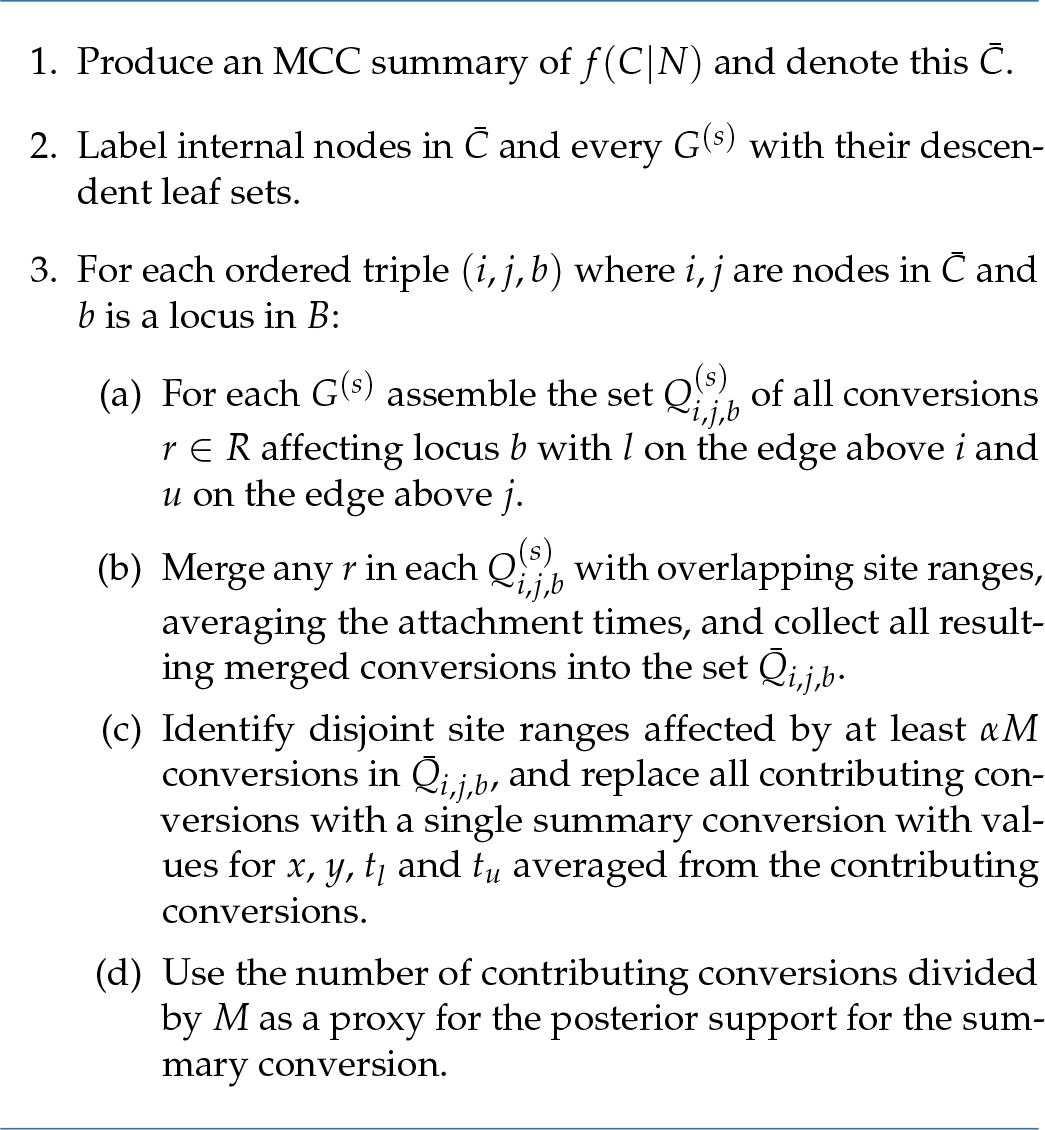
Method used to summarize samples *G*(^*s*^) for *s* ϵ [1,M] from the marginal posterior for *G*.

Testing with simulated data demonstrates that the method is capable of recovering useful summaries. However, one significant drawback is that the algorithm only groups together sampled conversions that appear between identical (in the sense described in the algorithm) pairs of CF edges. This means that a single conversion with significant uncertainty in either of its attachment points *u* or *l* may appear as multiple conversions in the summary. As a result, we still consider the problem of how best to summarize the posterior distribution over ARGs a target for future research.

## Implementation and validation

The methods described here are implemented as a BEAST 2 package. This allows the large number of substitution models, priors and other phylogenetic inference methods already present in BEAST 2 to be used with the ClonalOrigin model.

Despite the reuse of an existing phylogenetic toolkit, the implementation is still complex. As such, the importance of validating the implementation cannot be overstated. Our validation procedure involved two distinct phases: sampling from the ARG prior and performing inference of known parameter values from simulated data.

### Sampling from the ARG prior

This first phase of the validation involves using the MCMC algorithm to generate samples from *f_CO_*(*G*|*N*, *ρ*, *δ*), i.e. the prior distribution over ARG-space implied by the ClonalOrigin model. Unlike the full posterior density, we can also sample from this distribution via direct simulation of ARGs. Statistical comparisons between these two distributions should yield perfect agreement. Assuming that errors in both the MCMC algorithm implementation or the ARG simulation algorithm are unlikely to produce identically erroneous results, this is a stringent test of all aspects of our implementation besides calculation of the ARG likelihood.

**Figure 3.**
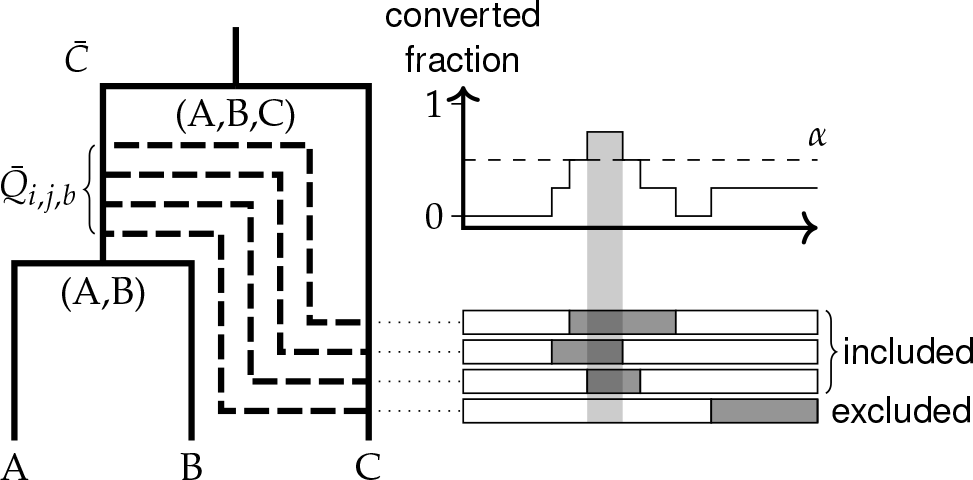
This diagram illustrates the way that conversions are summarized by algorithm 1. The solid tree on the left depicts the MCC summary of the clonal frame, *C*̄, with each node labeled by its set of descendent leaves. The dashed edges represent distinct conversions *Q*̄;_*i,j,b*_ that exist between a given pair of edges *i* and *j* in ARGs sampled from the posterior (with overlapping pairs of conversions persent on single ARGs merged). The horizontal boxes on the right indicate the site regions affected by each conversion, with the graph above showing the fraction of sampled ARGs posessing conversions at each site. A summary conversion is recorded only when this fraction exceeds the threshold *α*.

Figure 4 displays a comparison between the histograms for a number of summary statistics computed from ARGs with 5 (non-contemporaneous) leaves sampled using our implementation of each method. The MCMC chain was allowed to run for 10^8^ iterations with ARGs sampled every 10^4^ steps, while the simulation method was used to generate 10^5^ independent ARGs. The close agreement between the two sets of histograms is very strong evidence that our implementation of both algorithms is correct.

### Inference from simulated data

A common way to determine the validity and usefulness of an inference algorithm is to assess its ability to recover known truths from simulated data. In contrast with sampling from the prior, inference from simulated data is sensitive to the implementation of the ARG likelihood. We use here a well calibrated (Dawid 1982) form of the test, which requires that known true values fall within the estimated 95% highest posterior density (HPD) interval 95% of the time.

The details of the validation procedure are as follows. Firstly, 100 distinct 10-leaf ARGs of were simulated under the ClonalOrigin model with parameters *ρ* = 0.01, *δ* = 500 and *N* = 0.05. These ARGs were then used to produce an equivalent number of two-locus alignments, with each locus containing 5 ⨯ 10^3^ sites. Finally, each simulated alignment was used as the basis for inference of the ARG using the MCMC algorithm described above, conditional on the known true parameters.

The circles in the graphs shown as Figure 5 display the fraction of the sampled marginal MCMC posteriors for the CF tM-RCA (time to most recent common ancestor) and recombination event count which included the known true values as a function of the relative HPD interval width. The dashed lines indicate the fractions expected of a well-calibrated analysis. This close agreement therefore suggests that our analysis method is internally consistent in this regard, a result which strongly implies that our implementation is correct.

**Figure 4.**
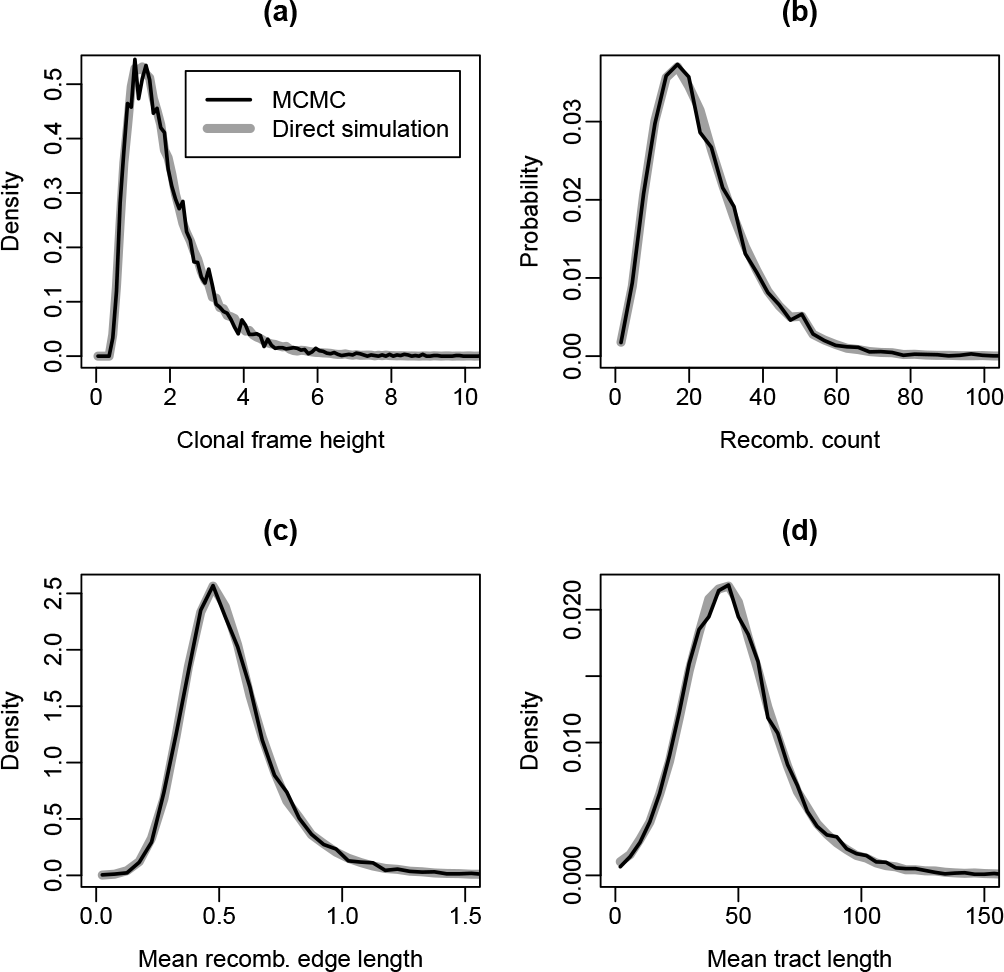
Comparison between distributions of summary statistics computed from ARGs simulated directly under the model (gray lines) and ARGs sampled using the MCMC algorithm (black lines). These include (a) the age of the CF root node, (b) the number of recombinations, and the average length of the recombinant (c) edges and (d) tracts on each sampled ARG. Exact agreement for each summary suggests that both algorithms are correct.

**Figure 5.**
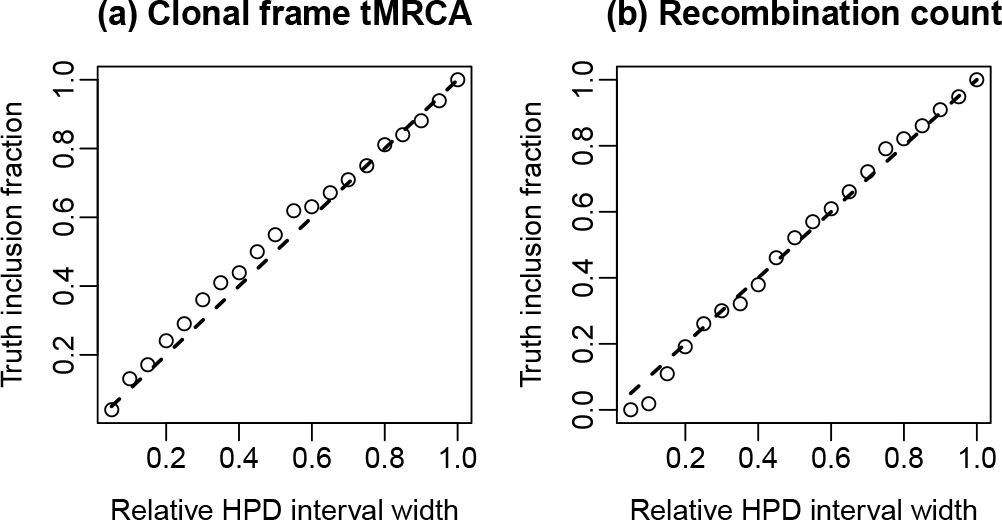
Coverage fraction versus HPD interval width for (a) the clonal frame tMRCA and (b) the recombination event count posteriors inferred from simulated sequence data. The circles represent the observed coverage fraction, while the dashed lines indicate the coverage fraction to be expected from a well-calibrated analysis.

## Example: Escherichia coli

We applied our new method to the analysis of sequence data collected from a set of 23 *E. coli* isolates. The isolates were derived from from both humans and cattle and include both STEC (Shiga toxin-producing *E. coli*) and non-STEC representatives of the O26 and O157 serotypes. The analysis focused on the 53 loci targeted by ribosomal multi-locus sequence typing (rMLST, Jolley *et al.* 2012).

The analysis was performed under the assumption of a constant population, the size of which was given a log normal prior ln*N*(0,2). The HKY substitution model (Hasegawa *et al.* 1985) was used, with uniform priors placed on the relative site frequencies and a log normal prior ln *N*(1,1.25) placed on the transition/transversion relative rate parameter *κ*. For the relative recombination rate *ρ*/*μ* we use an informative log normal prior of ln*N*(−2.3,1.5) which includes a previously published 95% credibility interval of 0.03–2.0 (Vos and Didelot 2009). The expected tract length parameter was fixed at *δ* = 10^3^ sites.

Six unique instances of the MCMC algorithm were run in parallel. Five of these were run for 2.5 ⨯ 10^7^ iterations while the sixth was run for 5 ⨯ 10^7^ iterations. Comparison of the posteriors sampled by each of these chains demonstrated that convergence had been achieved. Final results were obtained by removing the first 10% of samples from each chain to account for burn-in and then concatenating the results. Once complete, the effective sample size for every model parameter and summary ARG statistic recorded surpassed 200.

The final results of this analysis are presented as figure 6. Firstly, figure 6a displays a summary ARG produced from the sampled ARG posterior using a conversion posterior cutoff threshold of 0.4. This summary shows that four conversion events have posterior support exceeding this threshold. Three of these depict gene conversion events that transfer nucleotides between lineages ancestral to samples with O157 serotype. More specifically, the conversions result in gene flow from lineages ancestral to pathogenic (+STEC) samples to lineages ancestral to non-pathogenic (−STEC) samples. The remaining conversion event is indicative of a recent introgression from the O26 serotype into −STEC O157.

**Figure 6.**
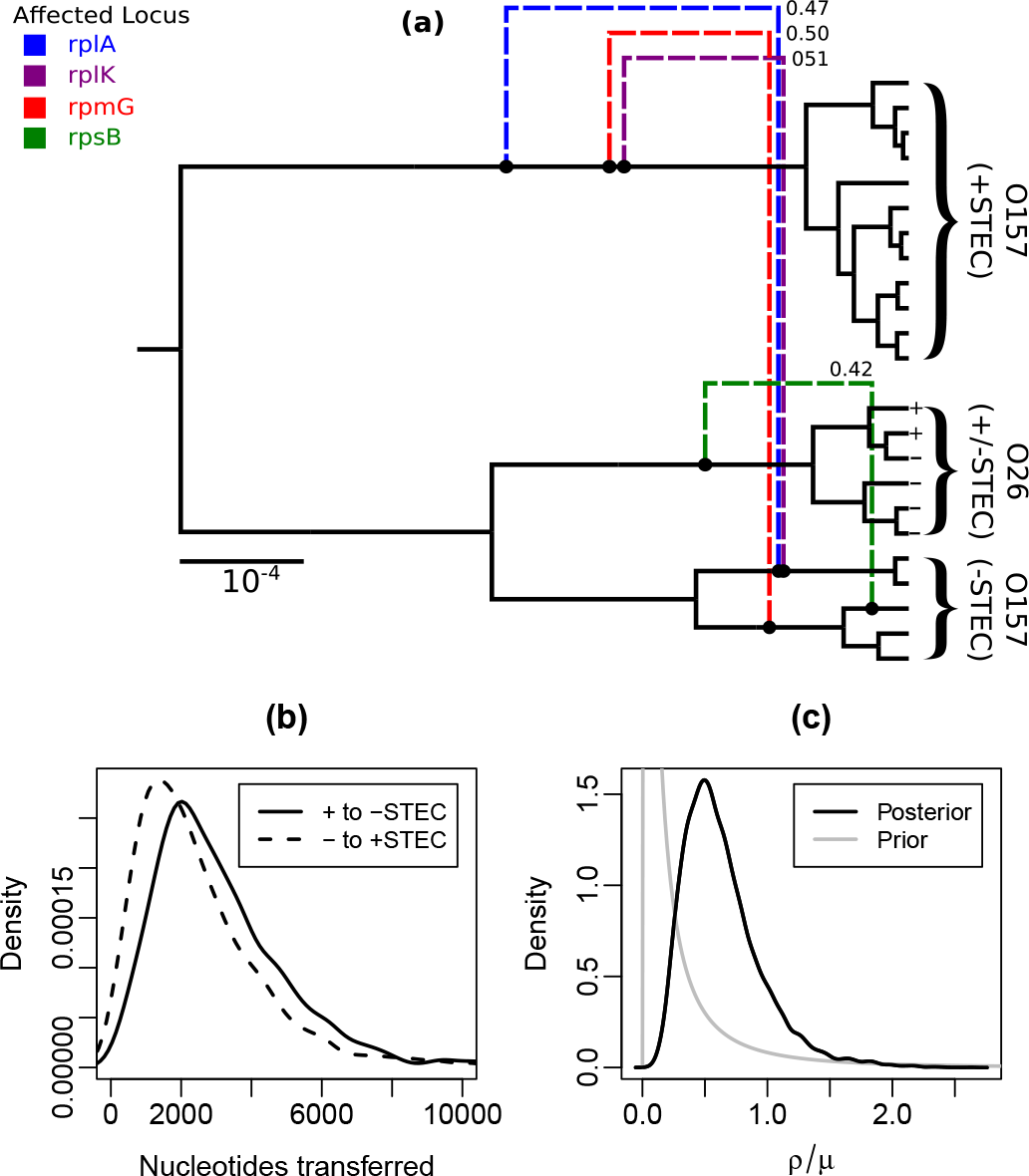
(a) Summary ARG produced by applying our method to sequences obtained from 23 *E. col; isolates*. Dashed edges represent summary conversions, with the numbers giving the estimated posterior support values. Conversions originating from the root edge of the clonal frame have been omitted.(b) Posterior distributions over nucleotides transferred between lineages ancestral to +STEC and -STEC O157 samples. (c) Posterior and prior distributions for the relative recombination rate, *ρ*/*μ*.

This overall pattern is also reflected in figure 6b which displays the posterior distributions for the total number of nucleotides transferred by conversion events between +/−STEC O157 ancestral lineages: the gene flow from +STEC to −STEC O157 is on average greater than that in the reverse direction. This asymmetry is, however, very slight—a fact which may be attributed to the presence of a large number of “background” conversions which individually lack the posterior support to be included in the summary but which nevertheless contribute to the particular gene flow metric we have chosen.

Finally, figure 6c displays the posterior distribution for the relative recombination rate parameter, giving a 95% HPD interval of [0.21,1.44]. The log-normal prior density for the recombination rate is also shown and indicates that the data are informative for this parameter.

The data and BEAST 2 XML files needed to replicate both this analysis and the validation studies are provided as part of the online supplementary material.

## Discussion

Dealing appropriately with recombination in a phylogenetic setting is a difficult task for a number of reasons. Firstly, the progressive bifurcation of lineages with increasing age steadily decrease the signal for these features in a given dataset. Furthermore, the possibility of these bifurcations drastically increases the size of the state space occupied by the genealogy. Indeed, even for a small number of aligned sequences, the upper bound of the number of coalescent events influencing the evolution of those sequences is potentially huge: the total number of nucleotide sites in the alignment. Considering that the super-exponential rate at which the number of binary trees grows as a function of sample size already presents complexity problems for computational phylogenetics, it is no surprise that models that explicitly consider recombination are not as widely used in genealogical inference.

Despite these challenges, Didelot and coauthors have shown repeatedly that traditional coalescent-based phylogenetic inference methods can be applied to such models, by applying carefully chosen simplifications to the coalescent with gene conversion which reduce the state space while maintaining sufficient realism in the important context of bacterial evolution. In our paper we have sought to continue in this tradition, and have demonstrated that one can indeed perform full joint inference of tree-based ARGs using a carefully constructed MCMC algorithm. In contrast with previous methods (as implemented in ClonalOrigin), this approach more accurately characterizes the posterior for the ARG and should yield more accurate estimates of statistical uncertainty.

Also, in our effort to narrow the technological gap between inference using the ClonalOrigin model and Bayesian inference performed using common non-recombination-aware models, we have introduced a means of summarizing sampled tree-based ARG posteriors that is reminiscent of the methods often employed to summarize sampled tree posteriors.

We must emphasize, however, that despite making significant headway we do not consider either the ClonalOrigin inference problem nor the problem of summarizing posterior distributions over tree-based networks to be in any way “solved”. In the case of the inference problem, computational challenges relating to the way the algorithm scales with increasing frequency of recombination remain. These relate most directly to the large computational complexity of the ARG likelihood (eq. (5)). (This is often the most computationally expensive calculation even in standard phylogenetic analyses, and recombination only multiplies this burden.) It may be the case that improving this situation will require replacing the mathematically exact likelihood evaluation under a given substitution model with a carefully chosen approximation, but the feasibility and usefulness of this approach has yet to be fully investigated.

The problem of summarizing posterior distributions over tree-based networks would seem to be a fruitful line of future research. The algorithm presented here does seem to perform relatively well from an empirical standpoint, and to our knowledge is the first of its kind. However, it does have drawbacks relating to its propensity to misclassify conversions for which topological uncertainty exists (i.e. uncertainty in the CF edge to which one or both of its end-points attach) as multiple distinct conversions with a proportionally smaller posterior support. Solving this problem would seem to be non-trivial, as it requires the algorithm to identify a conversion in one sampled ARG with a conversion in a second ARG even when those conversions join distinct pairs of edges on the clonal frame. However, we feel that tackling these and other related problems is a worthwhile endeavor and one which should encourage mainstream adoption of recombination-aware Bayesian phylogenetic inference methods.

## Appendix: MCMC state proposal distributions

In this appendix we lay out the details of the proposal operators used by the MCMC algorithm implemented described in the paper. To do this, we require some additional nomenclature. We decompose the CF using the tuple *C* = (*V*, *E*, **t**). Here *V* = *I* ⋃ *Y* with *I* being the set of leaf nodes and *Y* being the set of internal nodes, which contains the root node *o*. The set E contains the directed edges between nodes *i*, *j* ϵ *V*, where an edge from *i* to *j* is written 〈*i*, *j*〉. We use t = {*t_i_* |*i* ϵ *V*} to denote a set of node ages. The direction of an edge 〈*i*, *j*〉 is such that *t_i_* < *t_j_*.

As noted in the manuscript, MCMC is an iterative algorithm for sampling from some target probability density *π*(*x*) by iteratively modifying the state *x*. At each step in the iteration, a specific proposal kernel *q_w_*(*x*′|*x*) is chosen from a fixed weighted distribution of such kernels, and a new value for the state x^′^is drawn using that proposal. This new value is accepted with probability
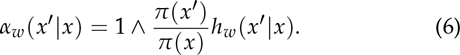
If the value is accepted it is assigned to *x*, otherwise *x* remains unchanged. The process then repeats. The term *h_w_*(*x*′|*x*) is a function which we refer to as the *Hastings-Green factor* or HGF for the proposal distribution, and ensures that the Markov chain defined by the MCMC algorithm is reversible. The HGF is uniquely defined by the proposal, but is often non-trivial to derive. Thus, each operator is presented below alongside its corresponding HGF.

## A ARG scale proposal

This operator selects a scaling factor *f* uniformly at random from [*α*, *α*^−1^] where *α* ϵ (0,1) is a tuning parameter for which smaller values yield bolder proposals. The age of every entity in the ARG, excluding leaf ages, is scaled by this one factor. The HGF for this proposal is
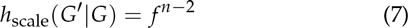
where *n* is the number of entities scaled by the move.

## B Conversion add/remove

With probability 1/2 this operator either deletes a randomly selected conversion, or creates a new conversion *r* = (*l, u, b, x, y*) drawn directly from the prior
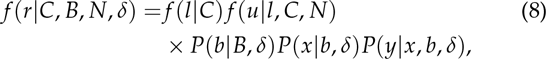
where the terms on the right-hand side are those described in the manuscript. The HGF for the deletion form of the proposal is
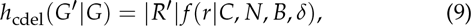
where *r* is the conversion selected for deletion.

The HGF for the addition form is simply *h*_cadd_(*G*′|*G*) = 1/*h*_cdel_ (*G*|*G*′).

## C. Detour add/remove

This operator improves mixing by allowing the sampler to transition directly between ARGs that have very similar local tree sets. It does this by proposing the addition or deletion of “detours“: pairs of conversions (*r*_1_, *r*_2_) for which *u*_1_ and *l*_2_ lie on the same edge of *C* and for which the attachment times satisfy *t_u1_* < *t_l2_*.

With probability 1/2 either the deletion or the addition form of the operator is selected. For addition, a conversion *r* is selected uniformly at random from *R*. Two times *t_d1_* and *t_d2_* are drawn from Unif(*t_l_*, *t_u_*) and labeled so that *t_d1_* < *t_d2_*. A non-root node *i* is then chosen uniformly at random from *V*. Let *i_p_* be the parent of *i*. If *u* or *l* lie on 〈*i*, *i_p_*〉 or it is not the case that both *t_d1_*, *t_d2_* ϵ [*t_i_*, *t_ip_*] then the proposal is immediately rejected. Otherwise, *r* is replaced with a pair of conversions *r*′_1_ = (*l*, *u*′, *b*, *x*, *y*) and *r*′_2_ = (*l*′, *u*, *b*, *x*′, *y*′) where *l*′ and *u*′ are the points on 〈*i*, *i_p_*〉 with times *t_d1_* and *t_d2_* respectively, and *x*′ and ′*y* and *b*′ are drawn from the affected site region boundary priors *P*(*b*|*B*, *δ*), *P*(*x*|*b*, *δ*) and *P*(*y*|*x*, *b*, *δ*).

For deletion, a non-root node *i* is chosen uniformly at random from *V*, and *i_p_* is defined as its parent. A pair of conversions *r*_1_ and r_2_ are chosen uniformly at random satisfying the requirements *u*_1_ ≠ *l*_1_, *u*_2_ ≠ *l*_2_, *u*_1_ lies on 〈*i*, *i_p_*〉 and *l*_2_ lies on 〈*i*,*_ip_*〉. This pair is replaced by a single conversion *r*′ = (*l*_1_,*u*_2_,*b*_1_,*x*_1_,*y*_1_). The HGF for the addition form is
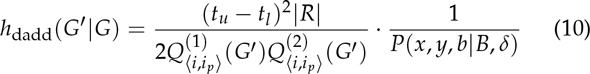
where 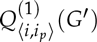 is the number of conversions *r*′′ in *R*′ where *u*′′ and *l*′′ lie on distinct CF edges and where *u*′′ lies on 〈*i*, *i_p_*}. Similarly 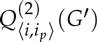 is the number of conversions with *u*′′ and *l*′′ on distinct edges and where *l*′′ lies on 〈*i*, *i_p_*〉. For the deletion form the HGF is 
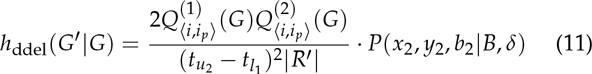

## D. Redundant conversion add/remove

This operator adds or removes a conversion that mirrors an existing edge in *C*, meaning that the conversion doesn′t introduce a change in the local tree topology. The boldness of the move is adjustable via the tuning parameter *λ*.

With probability 1/2 the addition or removal form of the operator is selected. For addition, a non-root node *i* is drawn uniformly at random from *V*, and *i_p_* is defined as its parent. A new conversion *r* = (*l*, *u*, *b*, *x*, *y*) is created with *x*, *y* and *b* drawn from the prior *P*(*x*, *y*, *b*| *B*, *δ*). The departure point *l* is drawn uniformly from the portions of edges around *i* with an age difference of at most *λ* from *t_i_*. Similarly, *u* is drawn from the portions of edges around *i_p_* that differ in age by at most *λ* from *t_ip_*.

For removal, a non-root node *i* is also drawn uniformly from *V*, with *i_p_* again defined as its parent. The sub-set 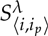 of *R* consisting of those conversions which could have been generated by the addition form of the move applied to the same CF edge 〈*i*, *i_p_*〉with a given *λ* is constructed. A member *r* of this set is selected uniformly at random and is deleted.

The HGF for the addition form is
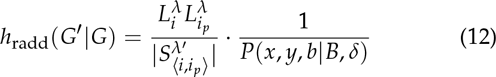
where 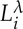 is the sum of the lengths of the CF edge portions around *i* from which *l* is drawn. Similarly, 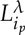 is the sum of the lengths of the CF edge portions around *i_p_* from which *u* is drawn. The primed 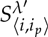 is the subset of *R*′ of conversions, including *r*, which could have been produced by this proposal.

For deletion, the HGF is
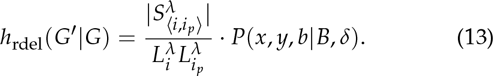

## E. Merge/Split conversion

This operator reversibly merges two conversions whose arrival and departure points share the same pair of CF edges.

A locus *b* is drawn from the prior *P*(*b*| *B*, *δ*). With probability 1/2 the merge or split form of the operator is selected. For merging, two conversions *r*_1_ and *r*_2_ are sampled without replacement from the subset *R*_*b*_ ⊂ *R* containing only those conversions affecting locus *b*. This pair of conversions is replaced by a new conversion *r*′ =(*l*_1_, *u*_1_, *b*, *x*_1_ ∨ *x*_2_,*y*_1_ ∧ *y*_2_).

For splitting, conversion *r* is drawn from *R*_*b*_. Let *i* be the CF node below the edge containing *l* and *j* be the CF node below the edge containing *u*, and define *i_p_* and *j_p_* to be the parents of these nodes (in the instance that *j* is the root CF node, *j_p_* is not defined). Two sites *m*_1_ and *m*_2_ are drawn uniformly from the site range [*x*, *y*]. With probability 1/2 we either define *x′_1_* = *x* and *x′_2_* = *m_1_* or *x′_1_* = *m_1_* and *x*′_2_ = *x*. Similarly, with probability 1/2 we either define *y*′_1_ = *y* and *y*′_2_ = *m*_2_ or *y*′_1_ = *m*_2_ and *y*′_2_ = *y*. Additionally, *l*′_2_ is a uniformly sampled point on the edge 〈*i*, *i_p_*〉. In the case that *j* is not the root, *u*′_2_, is sampled uniformly from 〈*j*, *j_p_*〉. Otherwise, the difference between the age of *u*′, *t*_*u*‱_, and the age of the root, *t*_j_, is drawn from the exponential distribution Exp (1/ (*t_u_ − tj*)). Conversion *r* is then replaced by a pair of conversions *r*′_1_ = (*l*, *u*, *b*, *x*′_1_, *y*′_1_) and *r*′_2_ = (*l*′_2_, *u*′_2_, *b*, *x*′_2_, *y*′_2_).

The HGF for the merge form is
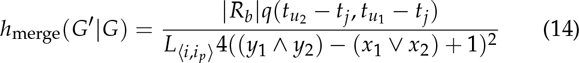
and for the split form is
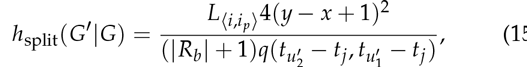
where 
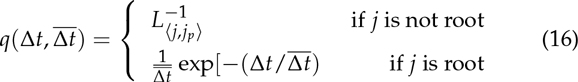

## F. Converted edge hop

This operator simply repositions the arrival or departure point of a randomly chosen conversion to be a new point on the tree. It proceeds by choosing a conversion *r* uniformly at random from *R*. Then, if *u* is above the root of *C* or with probability 1/2, *l*′ is drawn from a uniform density over *C* and *u*′ is set to *u*. Otherwise, *l*′ is set to *l* and *u*′ is drawn from a uniform density over *C*. In either case, if *t*_*u*′_ > *t_l_*, then *r* is replaced by a new conversion *r*′ = (*l*′, *u*′, *b*, *x*, *y*). If this condition is not met, the proposal is rejected.

The HGF for this move is unity.

## G. Converted edge flip

This is a simple proposal which reverses the direction of gene flow resulting from a given conversion. It is especially useful when this direction is not informed strongly (or at all) by the data. It involves firstly selecting a conversion *r* uniformly from *R* and defining *e_l_* as the CF edge containing the departure point *l* and *e_u_* as the CF edge containing the arrival point *u*. If *t_u_* falls outside of the time interval spanned by *e_l_* or *t_l_* falls outside of the time interval spanned by *e_u_* the proposal is immediately rejected. Otherwise, we then define new departure and arrival points *l*′ and *u*′ such that *t_l′_* = *t_l_* and *t_u′_* = *t_u_* but with *e_l′_* = *e_u_* and *e_u′_* = *e_l_*. Finally, we replace the conversion *r* with *r*′ = (*l*′, *u*′, *b*, *x*, *y*).

The HGF for this move is unity.

## H. Converted edge slide

This proposal ′slides′ a randomly selected arrival or departure point up or down the CF, with maximum size of the slide relative to the height of *C*, *t_o_*, is fixed by a tuning parameter *β* ϵ (0,1).

Firstly, the conversion is selected uniformly from *R* and a CF attachment point *p* is chosen uniformly from {*l*, *u*}. An age increment Δt is then drawn uniformly from [−*βt*_0_, *βt*_0_]. In the instance that Δ*t* > 1, the new attachment point *p*′ (i.e. *l*′ or *u*′ depending on the choice of *l* or *u* for *p*) is chosen to be that point on the lineage ancestral to *p* with *t_p′_* = *t_p_* + Δ*t* (If *p = l* and *t_p′_* > *t_u_* ∧ *t_o_* the move is immediately rejected.)

On the other hand, if Δ*t* < 0 the new attachment point *p*′ is chosen to be a point on a descendant lineage with *t_p′_* = *t_p_* + Δ*t*. (If *p* = *u* and *t_p′_* < *t_l_* the move is immediately rejected.) In the instance that *t*_*p*′_ is smaller than the age of the node below the CF edge containing *p*, there are multiple points on descendant lineages that satisfy this requirement. A particular point is chosen by tracing the CF lineage down from *p* and uniformly selecting the left or right child lineage of any CF node that is passed along the way to the final point *p*′. (If a leaf CF node is passed during this procedure the move is rejected immediately.)

In either case, the original conversion *r* is replaced by a new conversion *r*′ defined as either (*l*′, *u*, *b*, *x*, *y*) or (*l*, *u*′, *b*, *x*, *y*) depending on whether *p* represents an arrival or departure point, respectively.

The HGF for the move is 
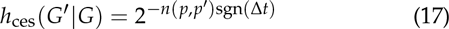
where *sgn*(Δ*t*) is the sign of Δ*t* and where *n*(*p*, *p*′) is the number of nodes on the CF on the lineage between points *p* and *p*′.

## I. Converted region swap

This proposal simply involves drawing two conversions *r*_1_ and *r*_2_ uniformly without replacement from *R* and swapping the loci and site ranges they affect. That is, the pair is replaced by a new pair *r*′_1_ = (*l*_1_, *u*_1_, *b*_2_, *x*_2_, *y*_2_) and *r*′_2_ = (*l*_2_, *u*_2_, *b*_1_, *x*_1_, *y*_1_).

The HGF for this move is unity.

## J. Converted region (boundary) shift

The converted region shift and converted region boundary shift propose adjustments to the region affected by a given conversion. Both use a tuning parameter *γ* that defines maximum size of the adjustment that can be made. The proposals begin by a conversion *r* is selected uniformly at random from *R*. A shift amount Δ is then drawn uniformly from [−*l_b_γ*/2, *l_b_γ*/2]. In the case of the region shift proposal, *x*′ = *x* + Δ and *y*′ = *y* + Δ. In the case of the region boundary shift proposal, either *x*′ = *x* + Δ and *y*′ = *y* or *x*′ = *x* and *y*′ = *y* + Δ with probability 1/2 The proposal is immediately rejected if either *x*′ or *y*′ lie outside of the allowed site range [1, *l_b_*] for locus *b*. The conversion *r* is then replaced by a new conversion *r*′ = (*l*, *u*, *b*, *x*′, *y*′).

The HGF for this move is unity.

## K. Clonal frame operators

With the exception of the topology-preserving temporal scaling operator, every move described thus far has proposed changes only to the set of conversions *R* applied to *C*, not *C* itself. Operators which propose changes to *C* are clearly of central importance to an algorithm designed to explore the joint (*R*, *C*) state space. As explained in the main text, our strategy for exploring this space is to employ each of the tree operators described in Drummond *et al.* (2002) to propose changes to *C*, updating *R* concurrently in order to maintain compatibility between the conversions and the CF. This is managed by expressing each of these operators primarily in terms of two primitive operations: *expand* and *collapse*. Understanding each operation requires considering a non-root node *i*, its parent *i_p_*, grandparent *i_g_* (if it exists) and sibling *i_s_* in *C*, as well as a distinct node *j* and its parent *j_p_* (if it exists) in *C* chosen so that *j* ∉ {*i*, *i_p_* } and *j* is not included in the subtree below *i*. Each operation involves “disconnecting” the subtree rooted by *i* from the rest of the clonal frame and “reconnecting” it to the edge above *j*. That is,
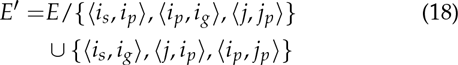
(Edges involving *j_p_* and *i_g_* are only included if these nodes exist.) This rearrangement is of course only valid if t is also updated so that 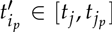 if *j_p_* exists or 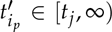 if it *j* is root in *C*. If such a modification is impossible, the proposal invoking the expansion or collapse is rejected immediately.

In terms of their effect on the CF, the only difference between the two operations is the sign of the difference 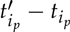: expansions increase the age of *i_p_* while collapses decrease this age. The effects on the set *R* of conversions are quite different, however.

For expansion, the set of conversion connections 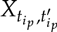 containing only those connections with 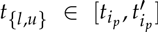 is constructed. Each of these attachment points are, with probability 1/2, moved in *R*′ to the contemporaneous point on the newly lengthened edge 〈*i*, *i_p_*〉. Additionally, in the case that *j* is the root of *C* (making *i_p_* the root of *C*), a set Z′ of new conversions are initiated along edges 〈*j*, *i_p_*〉 and 〈*i*, *i_p_*〉 with arrival points uniformly distributed amongst the portion of these edges at ages greater than *t_j_* ∨*t_ip_*. The expansion operation makes the following contribution to the HGF:
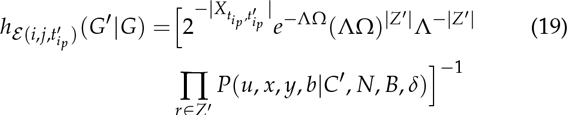
where 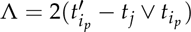 and Ω = Σ_*bϵB*_(*ρL*_*b*_ + *δ* − 1).

For collapse, the set of conversion connections 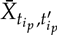 containing only those connections which lie on 〈*i*, *i_p_*〉 which have 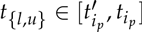 is constructed. Note that in the case that *i_p_* is the root of *C*, this set omits any attachment points belonging to conversions with arrival points 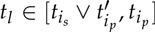. Such conversions are assigned to the set *Z*, along with conversions with arrival times in the same interval which lie on 〈*i_s_*, *i_p_*〉. Each attachment in 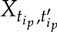 is moved to the lineage ancestral to *j*. Every conversion in *Z* is removed. The collapse operation makes the following contribution to the HGF:
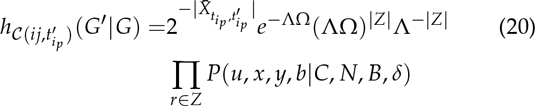
where 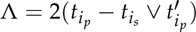 and Ω is as defined above.

We now describe each of the individual CF proposals. Note that with the exception of the CF/conversion swap operator (which is unique to our algorithm) we do not quantitatively describe how each move affects the CF, but instead explain how their operation is implemented in terms of expansions and contractions. Interested readers should refer to Drummond et al. (2002) to complete the descriptions.

**Uniform operator**: This operator proposes a new age 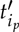 for randomly-selected non-root internal node *i_p_* within the interval imposed by the maximum age *t_i_* ∧ *t_is_* of its children, *i* and *i_s_*, and the age *t*_*i*_*g*__ of its parent, *i_g_*. This move is implemented as either a single expansion 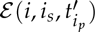 if 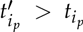 or a single collapse 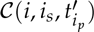 if 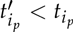.

**Subtree exchange operator**: This operator exchanges two distinct subtrees rooted by non-root nodes *i*^(1)^ and *i*^(2)^ and their respective parents, *i*_*p*_^(1)^ and *i*_*p*_^(2)^, and siblings 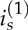 and 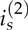. The operator is implemented via serial application of two primitive expand/collapse operations, with the type of operation determined by the relative ages of the parent nodes. If 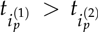 the operations are 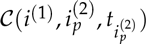 followed by followed by 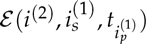. Otherwise, the operations are 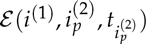 followed by 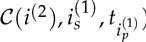.

**Wilson-Balding operator**: This operator takes a subtree rooted by the non-root node *i*, detaches it from the rest of the CF, then reattaches it to some other point at time 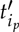 on the edge above a randomly chosen node *j*. (This is essentially the rooted time-tree equivalent of the Nearest Neighbour Interchange or NNI move used in walking the space of unrooted trees.) Besides selecting the nodes involved and the new time, this move involves just a single expand/collapse operation. If 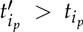 operation is 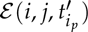, otherwise it is 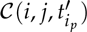.

**CF/conversion swap operator**: This final operator aims to in some sense swap the role of a conversion and a CF edge in describing a particular portion of the ARG topology. To do this, a conversion r is selected at random from the subset of *D* ⊆ *R* including only those conversions for which the arrival and departure points lie on distinct edges of *C*. The node below the edge containing, is labeled *i*, its sister *i_s_*, and the node below the edge containing *u* is labeled *j*. For the purpose of the expand/collapse operation, 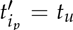. The conversion *r* is then replaced by *r*′ = (*l*, *u*′, *b*′, *x*′,*y*′), where *u*′ is the point on the edge above *i_s_* with time *t*_*i*_*p*__ and where *b*′, *x*′ and *y*′ define a new affected site range drawn from the prior *P*(*b*′, *x*′,*y*′|*B*, *δ*). Finally, if 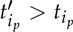 the expansion 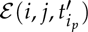 is performed, otherwise the collapse 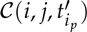 is performed. The HGF for this proposal is
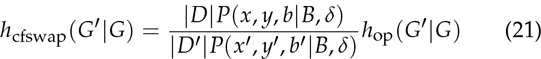
where *h*_op_ (*G*′|*G*) represents the HGF contribution of the particular expand/collapse operation performed.

